# LRP10 promotes trafficking of progranulin and prosaposin to lysosomes

**DOI:** 10.1101/2025.05.02.651888

**Authors:** Francesca Filippini, Swathi Devireddy, Shawn M. Ferguson

## Abstract

Mutations in LRP10, a low-density lipoprotein receptor family member, cause familial Parkinson’s disease and dementia with Lewy bodies. However, its direct cellular functions remain largely undefined. Using a multidisciplinary approach, our new data shows that LRP10 is required for the efficient trafficking of progranulin and prosaposin to lysosomes. Loss of LRP10 resulted in aberrant Golgi accumulations of progranulin and prosaposin and their reduced abundance in lysosomes. Disease-linked LRP10 missense mutations failed to support this lysosomal trafficking. Moreover, LRP10 KO mice developed striking microgliosis marked by enlarged and hyper-ramified microglia, accompanied by progranulin accumulation in the Golgi. Our results define LRP10 as a positive regulator of progranulin and prosaposin lysosomal protein trafficking and microglia homeostasis and thus shed new light on how its dysfunction may drive neurodegeneration in Parkinson’s Disease and dementia with Lewy bodies.

## Introduction

Progranulin (encoded by the GRN gene) is an important lysosomal protein (Zhou *et al*, 2019; Gillett *et al*, 2023; Root *et al*, 2024). Progranulin heterozygous loss-of-function leads to a neurodegenerative disease known as frontotemporal dementia (FTD) (Baker *et al*, 2006; Cruts *et al*, 2006; Gass *et al*, 2006) and homozygous loss causes neuronal ceroid lipofuscinosis, a lysosome storage disease (Smith *et al*, 2012; Tanaka *et al*, 2014). Human genetics studies have also linked GRN to risk for both Alzheimer’s disease and Parkinson’s disease (Cruchaga et al, 2013; Van Cauwenberghe et al, 2016; Wainberg et al, 2023; Ibanez et al, 2018; Nalls et al, 2021). Thus, based on human genetics and disease pathology, it is clear that progranulin contributes to normal lysosome functions and even modest reductions in progranulin abundance result in severe consequences (Cenik *et al*, 2012; Lee *et al*, 2017; Zhou *et al*, 2017). Within lysosomes, progranulin is cleaved into bioactive granulin peptides that play important roles in lysosomal homeostasis, although the field has yet to reach consensus on their exact functions (Cenik *et al*, 2012; Holler *et al*, 2017; Root *et al*, 2024; Hasan *et al*, 2023). Thus, not only is overall progranulin abundance important but so to is ensuring its efficient delivery to lysosomes. Progranulin trafficking relies on interactions with multiple proteins, including prosaposin, a key progranulin binding partner that links progranulin to receptors like Surf4 for ER to Golgi transport (Devireddy & Ferguson, 2022); the cation-independent mannose 6-phosphate receptor (CI-M6PR) for transport from the trans-Golgi network (TGN) to lysosomes and to LRP1 for endocytosis from the plasma membrane (Nicholson & Rademakers, 2016; Zhou *et al*, 2015a). Direct interactions between progranulin and sortilin at the plasma membrane and TGN additionally support the trafficking of progranulin to lysosomes (Hu *et al*, 2012; Carrasquillo *et al*, 2010). While these studies have made important progress towards defining mechanisms that support progranulin and prosaposin delivery to lysosomes, gaps remain in understanding of progranulin trafficking to lysosome. In particular, cell type specific contributions to ensuring efficient lysosomal delivery of progranulin and prosaposin in specialized cell types remain poorly understood.

In this study, we identify LRP10, a member of the low-density lipoprotein (LDL) family as a novel regulator of progranulin and prosaposin trafficking. Heterozygous mutations in LRP10 have been linked to Parkinson’s disease and dementia with Lewy bodies (Quadri *et al*, 2018; Li *et al*, 2021). LRP10 was previously reported to localize to cellular organelles including the TGN and early endosomes that are critical for trafficking newly made progranulin towards its final destination in lysosomes (Doray *et al*, 2008; Boucher *et al*, 2008b; Grochowska *et al*, 2021). The presence of acidic dileucine motifs in the cytoplasmic C-terminus of LRP10 haver furthermore been mechanistically linked to interactions with GGA proteins that promote sorting into vesicles that mediate trafficking between the TGN and endosomes (Boucher *et al*, 2008b; Doray *et al*, 2008). Building on our new bioinformatic analyses of gene expression that suggested a potential functional link between progranulin and LRP10, we observed that LRP10 is necessary for efficient delivery of progranulin and prosaposin to lysosomes in HeLa cells. In the mouse brain, progranulin trafficking in microglia was particulary sensitive to loss of LRP10. Furthermore, LRP10 mutations that cause Parkinson’s disease and dementia with Lewy bodies were unable to support progranulin trafficking. These discoveries establish LRP10 as a critical component of the trafficking machinery that sorts progranulin and prosaposin within the secretory pathway; reveals a previously uncharacterized role for LRP10 in supporting lysosomal function and demonstrates microglial vulnerability to LRP10 loss-of-function. Our findings thus advance understanding of lysosome cell biology and furthermore open new therapeutic avenues aimed at enhancing lysosomal protein delivery in neurodegenerative diseases.

## Results

### Identification of a link between LRP10 and progranulin

To investigate progranulin biology, we performed co-expression analysis using the SEEK platform (https://seek.princeton.edu/seek/) which ranks genes based on their expression similarity across multiple datasets (Zhu *et al*, 2015). This strategy is based on the prediction that genes that share related functions will often exhibit coordinated regulation of their expression. Strikingly, we found that prosaposin, a major known binding partner of progranulin, was a top hit when searching for genes whose mRNAs are co-expressed with progranulin (Figure 1A). There was also a major enrichment for other genes that encode lysosomes proteins (12 of the top 15 co-expressed genes).

**Figure 1.**
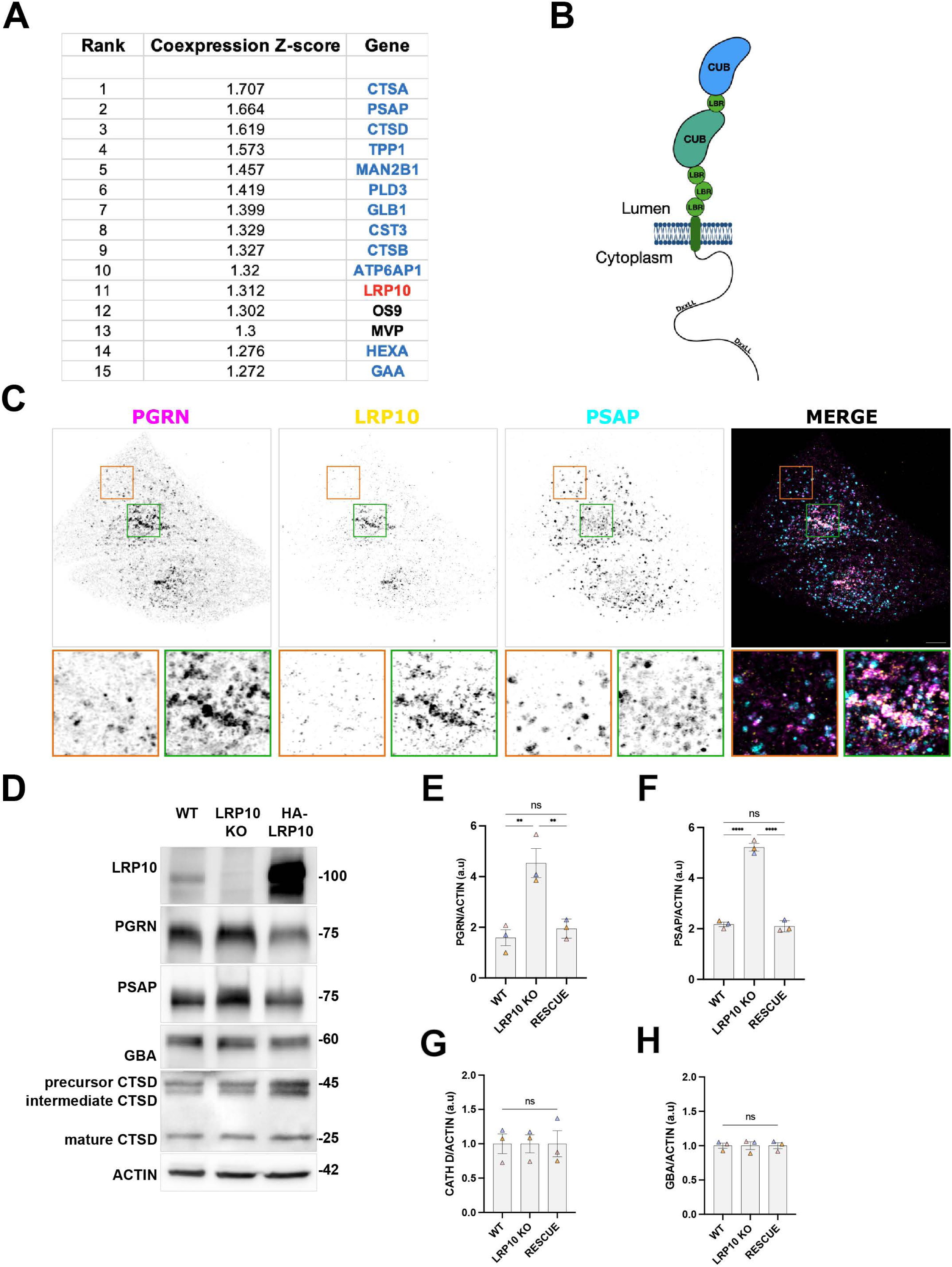
Identification of a key role for LRP10 in trafficking of progranulin and prosaposin. (A) Table showing results of PGRN co-expression analysis. Lysosomal proteins are displayed in blue. The strength of co-expression is represented by the Z-score, which indicates how many standard deviations a gene’s co-expression deviates from the mean background correlation. (B) Schematic diagram of LRP10 (left) showing two CUB domains in blue and green, four ligand binding-type domains (green spheres), a single transmembrane domain and acidic di-leucine sorting motifs (DXXLL) in the C-terminal region. (C) Immunofluorescence analysis of HeLa cells that stably express HA-LRP10 (yellow) were co-labeled for endogenous progranulin (PGRN, magenta) and prosaposin (PSAP, cyan). Projections of spinning disk confocal microscopy optical sections are shown. Scale bar, 5 μm. (D) Immunoblot analysis of the indicated proteins in wild-type, LRP10 knockout and HA-LRP10 rescued LRP10 KO cells (E-F-G-H). For quantification of immunoblot results, data from replicate experiments were Z-standardized by substracting the sample mean from each value and dividing by the sample standard deviation, resulting in values representing the number of standard deviations from the mean. To avoid negative numbers, the lowest value of all replicates was subtracted from the totality of values in the dataset. In each panel, mean (triangles) and SEM from 3 independent experiments are displayed. One-way ANOVA with Tukey’s multiple-comparison test is labeled on graphs. ∗∗p < 0.01; ∗∗∗∗p < 0.0001; ns non significant.

Gene ontology analysis of the top 100 genes whose expression was most similar to progranulin furthermore revealed a major enrichment for genes of the endolysosomal pathway (Figure S1A). This co-expression pattern between progranulin, prosaposin and numerous genes that encode lysosome proteins is consistent with well-established lysosome functions for progranulin and the close functional relationship between progranulin and prosaposin(Devireddy & Ferguson, 2022; Tran *et al*, 2023; Du *et al*, 2022; Paushter *et al*, 2018; Nicholson *et al*, 2016; Zhou *et al*, 2015b). Also, near the top of the list of genes co-expressed with progranulin was LRP10. A reciprocal SEEK search with LRP10 as the input revealed GRN as the number 2 hit and a similar overall endolysosomal pathway enrichment within the top 100 genes (Figure S1B). Although LRP10 had not previously been connected to progranulin or lysosomes, the combination between its expression pattern, reported localization to TGN and endosomes, domain organization and its membership in the LDL receptor family suggested a potential role for LRP10 in regulating the trafficking of progranulin, prosaposin and potentially other lysosome proteins (Doray *et al*, 2008; Boucher *et al*, 2008a; Grochowska *et al*, 2021; Pohlkamp *et al*, 2017)(Figure 1B).

Previous immunofluorescence assays and proteomic analyses of purified lysosomes support localization of LRP10 to the secretory pathway as well as the endolysosomal system (Boucher *et al*, 2008b; Doray *et al*, 2008; Brodeur *et al*, 2012; Grochowska *et al*, 2021; Akter *et al*, 2023). Consistent with these expectations, our immunofluorescence analysis of LRP10 showed that it strongly co-distributes with progranulin (but to a lesser degree with prosaposin) in the perinuclear region of Hela cells (Figure 1C). This partial overlap of LRP10 with each of these proteins is consistent with a putative function in controlling trafficking of these specific cargos through the secretory pathway. To test for a functional role for LRP10 in regulating the trafficking of progranulin, we used CRISPR-based genome editing to generate LRP10 KO HeLa cells. Immunoblot analysis of LRP10 KO cells revealed an accumulation of full length progranulin and prosaposin proteins (Figure 1D-F), but no change was detected for Cathepsin D or glucocerebrosidase (GBA, Figure 1D, G-H). The increased abundance of progranulin and prosaposin in LRP10 KO cells is consistent with selectively impaired delivery of progranulin and prosaposin to lysosomes and thus an accumulation of the full length, unprocessed, forms of these proteins. This accumulation of progranulin and prosaposin in LRP10 KO cells was alleviated upon re-expression of LRP10 (Figure 1D-F). These observations, together with the co-expression data, led us to hypothesize a critical role for LRP10 in the regulation of progranulin and prosaposin trafficking of progranulin and prosaposin from the Golgi to lysosomes.

### Impact of LRP10 knockout on progranulin and prosaposin localization

In the absence of LRP10, the progranulin signal that co-localizes with LAMP1-positive lysosomes (LAMP1) was significantly reduced and this was rescued by stable re-expression of HA-tagged LRP10 (Figure 2A). Consistent with an interaction between progranulin and prosaposin as they traffic through the secretory pathway (Devireddy & Ferguson, 2022), prosaposin localization to lysosomes was similarly affected by LRP10 KO and rescue (Figure 2C+D). In contrast, the localization of cathepsin D to lysosomes was unaffected by the absence of LRP10 (Figure 2E+F). In addition to their loss of lysosome localization, both progranulin and prosaposin clustered more in the perinuclear region of LRP10 KO cells (Figure 2A+C). Moreover, the progranulin lysosome localization was stronger in the HA-LRP10 rescued cells compared to WT, most likely due to over-expression of LRP10 in the rescue line (Figure 1D). These observations collectively support a role for LRP10 in the trafficking of progranulin and prosaposin to lysosomes.

**Figure 2.**
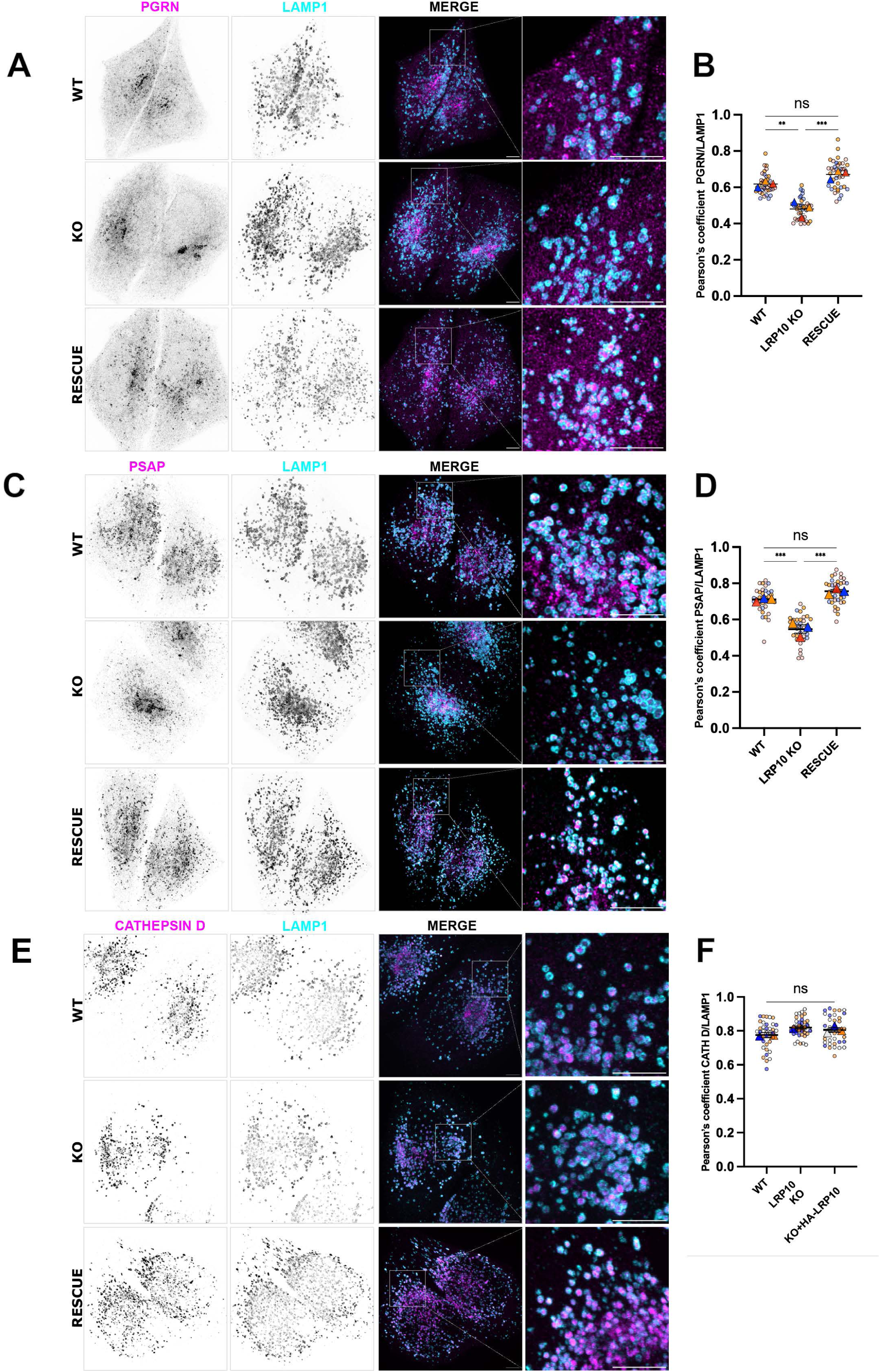
Impact of LRP10 knockout on progranulin and prosaposin localization. (A-B-C) Wild-type, LRP10 knock-out and stably expressing HA-LRP10 Hela cells were processed via immunohistochemistry and stained for LAMP1 (cyan) together with either progranulin (A, magenta), prosaposin (B, magenta) or cathepsin D (C, magenta) in C. Scale bar, 5 μm. (D-E-F) Superplots summarizing Pearson’s coefficient analysis of LAMP1 co-localization with the indicated proteins of interest. Horizontal lines show mean ± SEM, while colored circles and triangles represent values of individual cells and mean of 3 independent experiments (min 10 cells/experiment, ∗∗p < 0.01; ∗∗∗p < 0.001; ns non significant).

### Analysis of progranulin and prosaposin levels and processing within lysosomes from LRP10 KO cells

As an independent test of progranulin and prosaposin trafficking to lysosomes, we took advantage of an established lysosome purification assay based on endocytic pulse and chase of HeLa cells with superparamagnetic iron oxide nanoparticles (SPIONs) followed by cell rupture and magnetic capture of lysosomes (Figure 3A) (Amick *et al*, 2018). As seen in Figure 1, full length progranulin and prosaposin accumulated in whole cell lysates in the LRP10 KO cells (Figure 3B). Consistent with its known processing into granulins within lysosomes (Holler *et al*, 2017), full length progranulin was not readily detectable in lysosomes (Figure 3B and C). However, granulins were present in lysosomes but less abundant in the LRP10 KO samples. Likewise, both full length prosaposin and the saposin C fragment were less abundant in lysosomes from LRP10 KO cells (Figure 3B, D and E). We confirmed the specificity of the impact of LRP10 depletion on the trafficking of progranulin and prosaposin with the observation that there was no change in the abundance of mature cathepsin D in either LRP10 KO cell lysates or lysosomes (Figure 3B, F and G).

**Figure 3.**
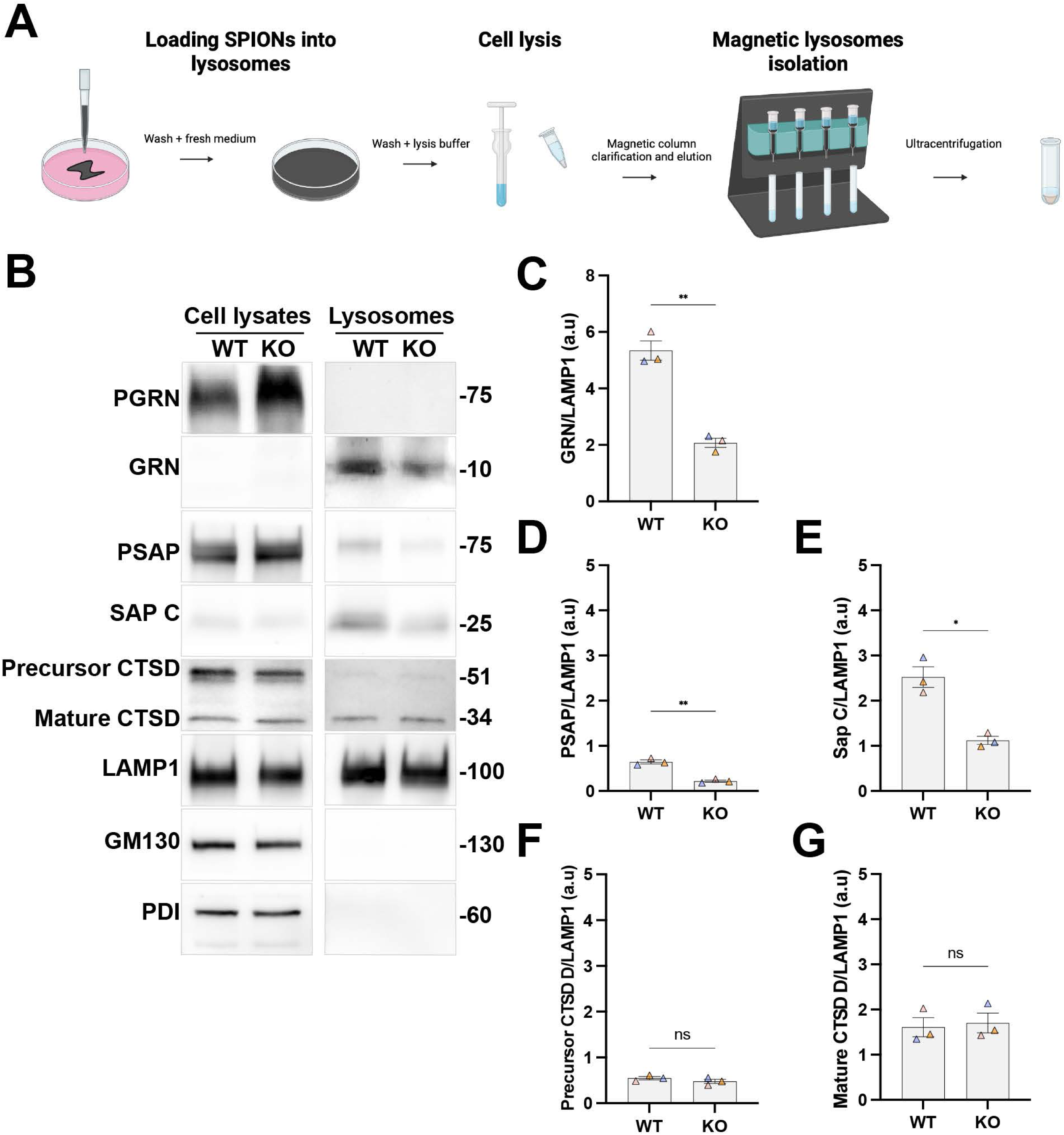
Depletion of granulins and saposin C from LRP10 KO lysosomes. (A) Schematic of lysosome isolation with superparamagnetic iron oxide nanoparticles (SPIONs, created in https://BioRender.com). (B) Immunoblot analysis of wild-type and LRP10 KO cell lysates and SPION-purified lysosomes. Direct comparison of organelle markers (LAMP1, GM130 and PDI) using same exposures. (C-D-E-F-G). Results for the cell lysate and lysosome fractions were derived from the same membranes. Lysates and lysosomes were loaded into a ratio of 10:1, respectively. For quantification of immunoblotting results, data from different replicate experiments were Z-standardized as described in the legend for Figure 3. In each panel, mean (triangles) and SEM from 3 independent experiments are displayed. T-test with Welch’s correction is labeled on graphs (∗p < 0.05; ∗∗p < 0.01; ns non significant).

### Progranulin and prosaposin are trafficked from the Golgi in a LRP10-dependent manner

To identify more precisely the progranulin and prosaposin trafficking step that is regulated by LRP10, we took advantage of the retention using selective hooks (RUSH) strategy that we previously established for characterization of Surf4-dependent ER-Golgi trafficking of progranulin and prosaposin (Devireddy & Ferguson, 2022). This strategy allows for the retention of proteins of interest in the ER and their synchronized release into the secretory pathway upon addition of biotin to the cell media. Before biotin addition, RUSH-tagged progranulin and prosaposin both showed the expected retention in the ER (Figure 4A-C). Approximately 20 minutes after biotin addition, both proteins accumulated at the Golgi apparatus before dispersing into vesicles that reflect delivery to into the endolysosomal pathway (Figures S2 and 4A-D) (Devireddy & Ferguson, 2022). Strikingly, we observed that in LRP10 KO cells both progranulin and prosaposin proteins failed to efficiently exit the Golgi by 40 min after biotin addition (Figure 4E-H). Collectively, these results point to a critical role for LRP10 in promoting trafficking of progranulin and prosaposin out of the Golgi.

**Figure 4.**
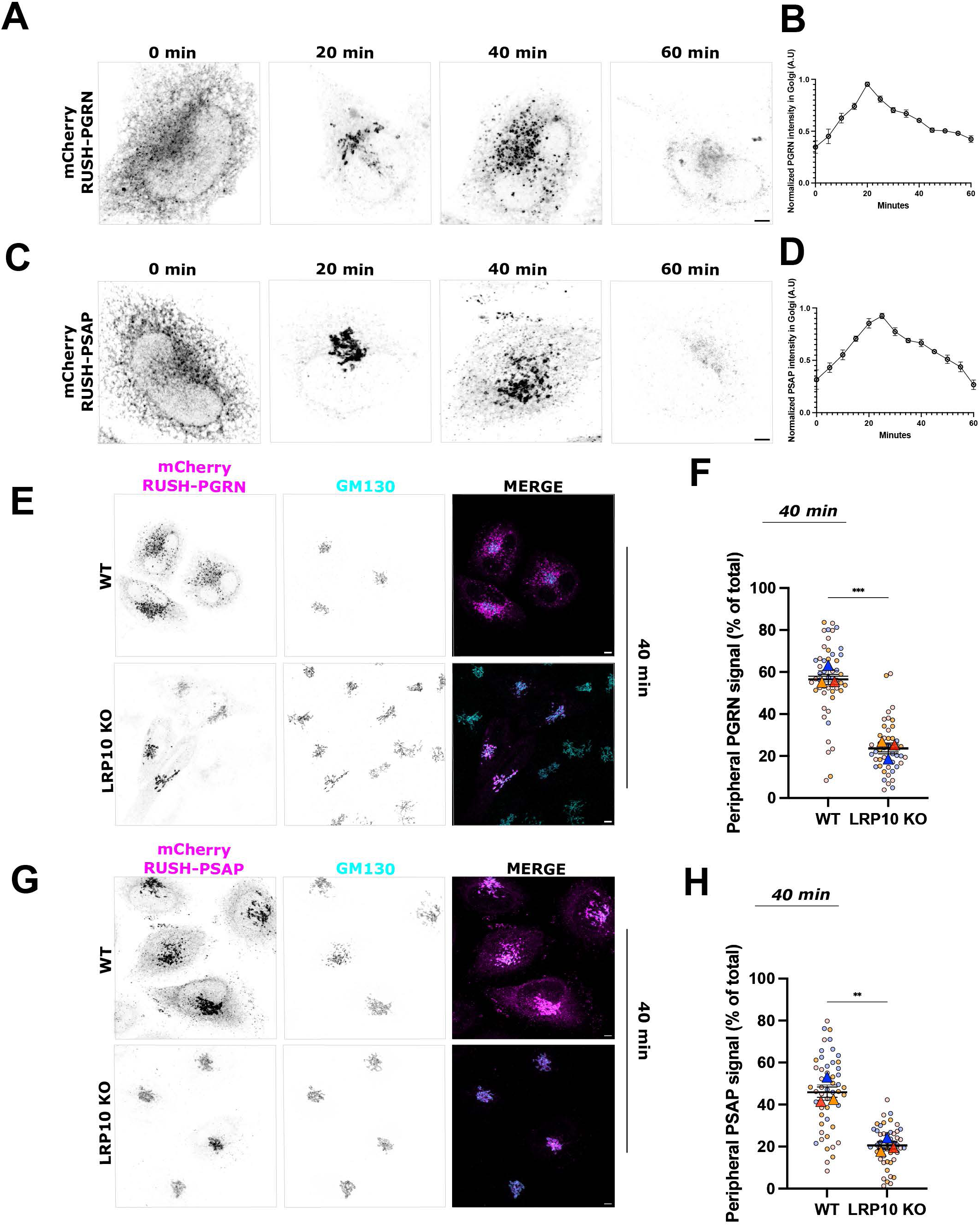
LRP10 promotes the efficient efflux of PGRN and PSAP from the Golgi. (A) Spinning disk confocal imaging of mCherry-RUSH-PGRN expressed in wild-type Hela cells at t = 0, 20, 40 min and 60 min after the addition of 40 µM biotin. (B) Tracking of mCherry-PGRN-RUSH protein intensity in the Golgi apparatus (via GM130 colocalization) through time. (C) Spinning disk confocal imaging of mCherry-RUSH-PSAP expressed in wild-type Hela cells at t = 0, 20, 40 min and 60 min after the addition of 40 µM biotin. (D) Tracking of mCherry-PSAP-RUSH protein intensity in the Golgi apparatus (via GM130 colocalization) through time. (E-F) mCherry-RUSH-PGRN and mCherry-PSAP-RUSH localization in WT and LRP10 KO Hela cells after 40 min of biotin incubation. Cells were immunostained for mCherry (magenta) and GM130 (Golgi protein, cyan). Z projections of confocal microscopy optical sections are shown. Scale bar, 5 μm. (G-H) Quantification of the mCherry-RUSH-PGRN and mCherry-PSAP-RUSH signal present at cell periphery (outside GM130 mask) over total signal, 40 min after biotin addition. Horizontal lines show mean ± SEM, while colored circles and triangles represent values of individual cells and mean of 3 independent experiments with at least 13 cells/experiment. T-test with Welch’s correction is labeled on graphs (∗∗p < 0.01; ∗∗∗p < 0.001).

### Progranulin Accumulation and microgliosis in LRP10 KO Mouse Brains

To investigate in vivo functions of LRP10, we generated a new strain of LRP10 KO mice (Figure S3). KO mice obtained from heterozygote matings were viable and reached adulthood.These matings yielded the expected 1:2:1 ratio (26 WT:45 Het.:23 KO, Chi squared test p=0.84) of WT, heterozygous and KO offspring that reached weaning age. This argues against loss of any KO mice due to either embryonic or perinatal lethality. We furthermore did not observe any overt health defects in the adult LRP10 KO mice. Given the role that we have uncovered for LRP10 as a regulator of progranulin and prosaposin trafficking and links of all these proteins to neurological diseases, we next looked at the impact of the LRP10 KO on the subcellular distribution of progranulin and prosaposin in the wildtype versus LRP10 KO mouse brains. Investigation of the localization of progranulin in the motor cortex of LRP10 WT and KO mouse brains revealed a punctate pattern for progranulin in neurons and glia that fits with the expectation of lysosome localization (Figure 5A). However, in the LRP10 KO, progranulin additionally accumulated in large clusters within microglia (Figure 5A-5B). Meanwhile, cathepsin D also exhibited a punctate localization pattern in the WT and KO brains but did not accumulate into large puncta in the soma of LRP10 KO microglia (Figure 5C). A modest progranulin accumulation pattern was also detectable in neuronal cell bodies, but much less strikingly than in microglia (Figure 5D). Analysis of LRP10 KO brains furthermore revealed evidence of microgliosis as shown by increased number of IBA1-positive microglia (Figure 5E). The microglia in LRP10 KO mice also exhibited larger somas and increased ramification compared to their WT counterparts (Figure 5A, E-H). This suggests an altered state of microglia in the absence of LRP10. Based on the known occurrence of microgliosis in GRN KO mice, dysregulation of progranulin trafficking in the absence of LRP10 may contribute to the microglia phenotypes arising from LRP10 KO (Wu *et al*, 2021). These in vivo findings in LRP10 KO mice extend our cell culture results in highlighting a crucial role of LRP10 in regulating progranulin trafficking and furthermore highlight a vulnerability of microglia to loss of LRP10 expression.

**Figure 5.**
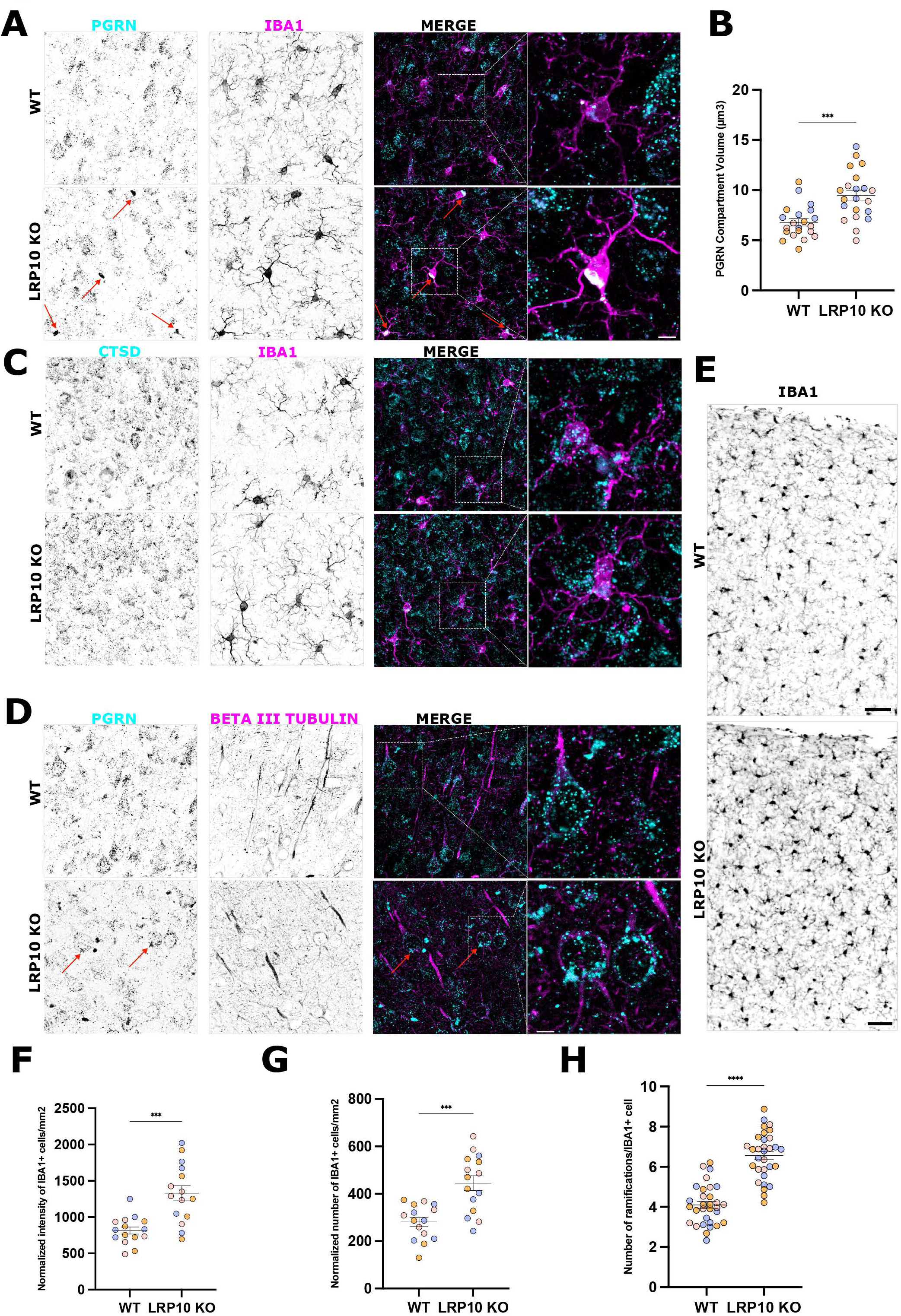
Microglia are vulnerable to the in vivo loss of LRP10. (A) Immunofluorescence analysis of the motor cortex region of brain sections from wild-type and LRP10 knock-out mice labelled for PGRN (cyan) and IBA1 (magenta). Mice were 13 months old and we analyzed sex matched pairs of WT and LRP10 KO mice across experimental replicates. Z projections of confocal microscopy optical sections are shown. (B) Quantification of PGRN signal in IBA1 positive cells of wildtype and LRP10 KO mice. IBA1 positive cells were manually selected from confocal Z-stacks and converted to 3D ROI in Icy (de Chaumont *et al*, 2012). The progranulin signal was segmented using spot detector (https://icy.bioimageanalysis.org/plugin/spot-detector/) and size of progranulin positive compartments was measured. Seven to ten cells per region of n = 3 pairs of mice were analyzed. Colors correspond to data from individual mice. T-test with Welch’s correction was performed from each triplicate mean values (∗∗p < 0.01). (C) PGRN (cyan) and Beta-III Tubulin (magenta) in WT and LRP10 KO motor cortex. (D) CTSD (cyan) and IBA1 (magenta) in WT and LRP10 KO motor cortex. (E) IBA1 staining in wild-type and LRP10 knock-out mice. Z projections of confocal microscopy optical sections are shown. Scale bar, 10 μm. (F) Quantification of number of IBA1 positive cells/mm^2^ using analyze particles in Fiji (Schindelin *et al*, 2012) and normalizing to the mean cell size in control samples. (G) Quantification of intensity of IBA1 positive cells/mm^2^ measuring the mean fluorescence intensity in Fiji and normalizing intensity values by the area (number of pixels in selected ROI). (H) Quantification of the total number of ramifications in IBA1 positive cells using Sholl analysis. 75-100 cells per region of interest from of n = 3 mice/genotype were used, colors correspond to individual mice. Scale bar = 5 μm in panels A, C and D; 10 µm in E.

### Progranulin accumulates in the trans-Golgi network in microglia of LRP10 KO mice

We further investigated the subcellular compartment where progranulin accumulates in microglia. We found that progranulin localizes to the TGN to a limited degree in WT mice and that it’s abundance at the TGN increased in the LRP10 KO microglia (Figure 6A). Interestingly, the overall size of the TGN was also larger in the LRP10 KO, potentially reflecting a traffic jam arising from the loss of LRP10-dependent cargo efflux (Figure 6A). By immunostaining brain sections with progranulin and prosaposin, we confirmed that these two proteins both co-accumulated within the same TGN clusters in the LRP10 KO microglia, although this effect was more striking for progranulin (Figure 6B).

**Figure 6.**
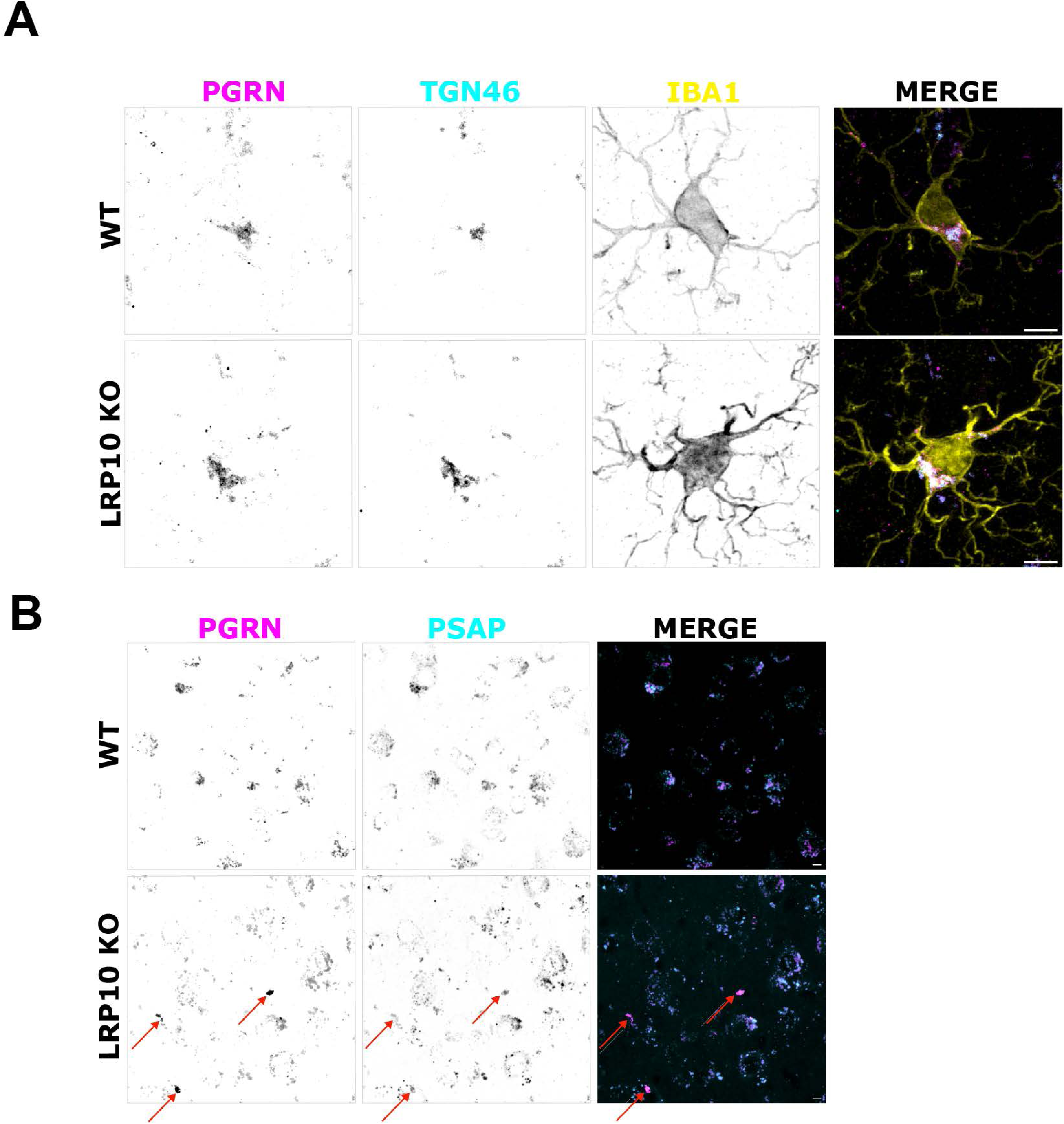
Progranulin and prosaposin accumulate at the trans-Golgi network in LRP10 KO microglia. (A) Brain sections from wild-type and LRP10 knock-out mice were labelled for PGRN (cyan), TGN46 (yellow) and IBA1 (magenta). Whole Z projections of confocal microscopy optical sections from the motor cortex are shown. Scale bar, 5 μm. (B) Brain sections from wild-type and LRP10 knock-out mice were labelled for progranulin (PGRN, magenta) and prosaposin (PSAP, cyan). Whole Z projections of confocal scanning microscopy optical sections from the motor cortex are shown. Scale bar, 5 μm.

### Parkinson’s Disease-associated LRP10 mutations prevent efficient delivery of progranulin to lysosomes

As LRP10 variants that lead to premature stop codons and missense mutations have been identified as causes of familial PD and LBD (Quadri *et al*, 2018), we investigated their impact on the transport of lysosomal proteins. Figure 7A presents an AlphaFold structural prediction, highlighting the locations of disease-causing mutations while Figure 7B provides a schematic diagram illustrating locations of LRP10 mutations. We tested the ability of these LRP10 mutants to rescue the lysosome localization of progranulin in LRP10 KO HeLa cells. Compared to WT LRP10 which robustly increased the lysosome abundance of progranulin, none of the LRP10 mutants were functional in this assay even though they all were expressed and had punctate subcellular localization consistent with an ability to fold and exit the ER (Figure 7C and S4).

**Figure 7.**
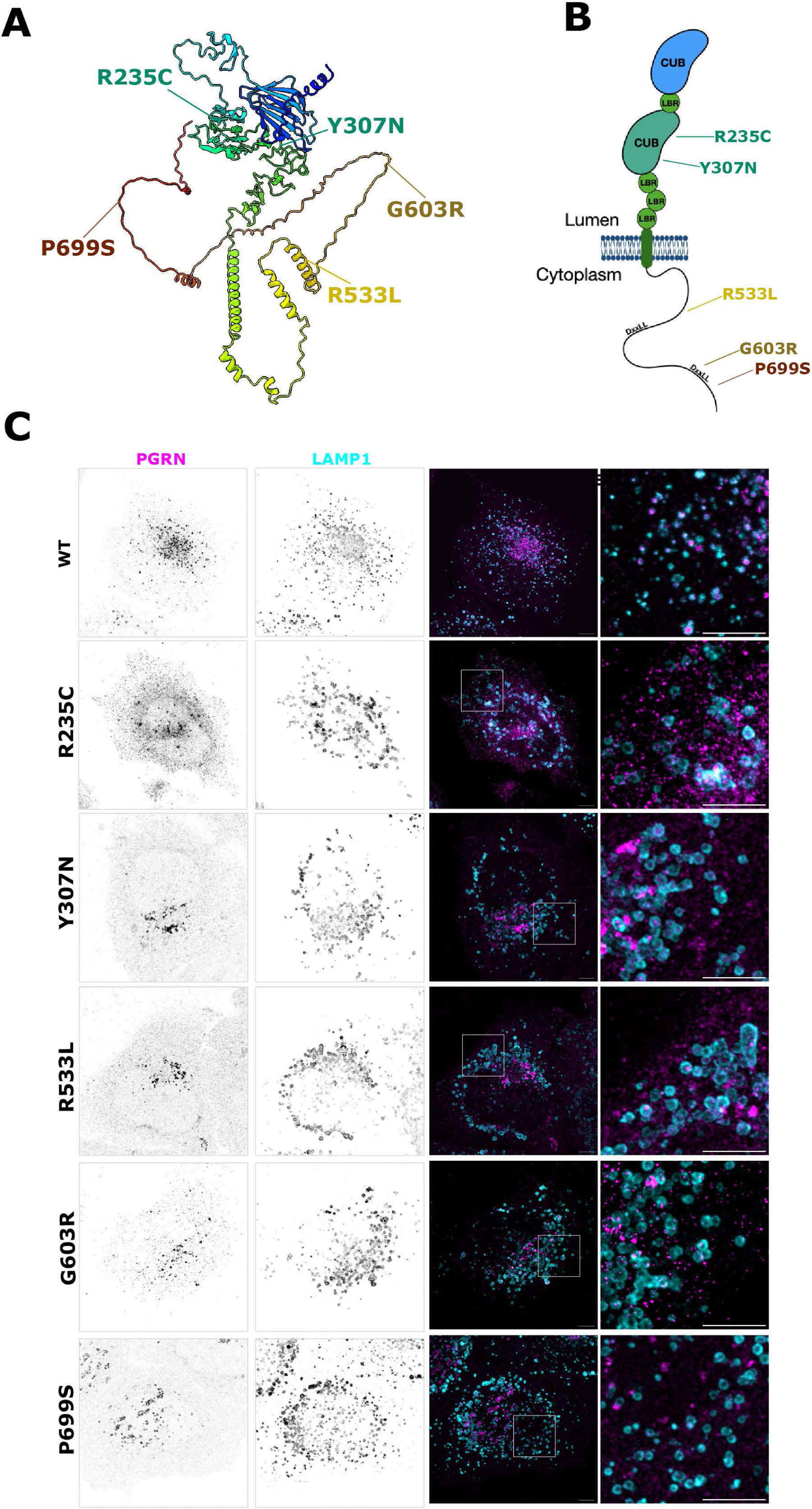
LRP10 Parkinson’s disease and Lewy body dementia mutants do not support PGRN trafficking to lysosomes. (A) Alphafold LRP10 structure prediction with PD/LBD associated mutations. (B) Schematic diagram of LRP10 domain organization with annotation of PD/LBD-associated mutations annotated. (C) LRP10 knockout Hela cells were transfected with wildtype or the indicated mutant HA-LRP10 plasmids and processed for immunofluorescent detection of PGRN (magenta) and LAMP1 (cyan). Scale bar, 5 μm.

## Discussion

In this study, we demonstrate that LRP10 contributes to the lysosomal trafficking of progranulin and prosaposin. LRP10 deficiency leads to disrupted lysosomal delivery of progranulin and prosaposin, thus implicating LRP10 in the regulation of lysosome homeostasis. Furthermore, we provide evidence that the absence of LRP10 results in microglia-specific alterations, characterized by microgliosis and the accumulation of progranulin and prosaposin within the Golgi. These findings indicate an important role for LRP10 in microglia and that loss of LRP10 may contribute to neuroinflammation and neurodegeneration due to defects the trafficking of progranulin and prosaposin to microglial lysosomes.

LRP10 shares functional similarities with known lysosomal sorting receptors, such as sortilin and the cation independent mannose-6-phosphate receptor (M6PR), both of which mediate the trafficking of PGRN and PSAP. Sortilin has been well characterized as a key regulator of progranulin transport to lysosomes, and M6PR is essential for trafficking of the progranulin-prosaposin complex (Zhou *et al*, 2015a, 2017). A simple explanation for the progranulin and prosaposin trafficking defects arising from LRP10 depletion is that a direct interaction between either progranulin and/or prosaposin with LRP10 allows LRP10 to sort progranulin-prosaposin into vesicles at the TGN that are destined for the endolysosomal pathway. However, we have not yet been able to demonstrate such an interaction between LRP10 and either PGRN or PSAP via immunoprecipitation. These negative results due not conclusively rule out the possibility of a direct interaction that is either of modest affinity or which was not preserved under our experimental conditions. As a result, we have focused this manuscript on the functional impact of LRP10 depletion on the trafficking of progranulin and prosaposin.

The microglia phenotype observed in LRP10 knockout (KO) mice provides new insight into a role of LRP10 in neuroimmune function. We observed that loss of LRP10 resulted in an increase in microglia abundance and process ramification, suggestive of an activated microglia state. Notably, these microglia exhibited marked Golgi-localized accumulation of PGRN, suggesting a failure in its trafficking to lysosomes. Given that PGRN is essential for microglial function and lysosomal integrity, its mislocalization in LRP10-deficient microglia could contribute to neuroinflammation and disease pathology (Lui *et al*, 2016)(Wu *et al*, 2021).

In addition to GRN-linked frontotemporal dementia, our findings have potential implications for Parkinson’s disease (PD) and dementia with Lewy bodies (DLB), both of which have been linked to LRP10 mutations (Vergouw *et al*, 2019; Gagliardi *et al*, 2021). Lysosomal dysfunction is increasingly recognized as a key contributor to PD pathogenesis (Dehay *et al*, 2013; Navarro-Romero *et al*, 2020; Bentley-DeSousa *et al*, 2025), and our results suggest that LRP10 mutations may confer disease risk by impairing lysosomal abundance of progranulin in microglia. At minimum, the emergence of both cell biological data that places progranulin and LRP10 in a common pathway along with human genetics data that link both GRN and LRP10 to Parkinson’s disease warrants further investigation.

Collectively, our results define a new role for LRP10 in promoting the trafficking of progranulin and prosaposin and demonstrate that microglia are particularly vulnerable to LRP10 deficiency. Future studies should focus on elucidating the molecular mechanisms underlying LRP10-mediated trafficking of progranulin and prosaposin and the basis for microglia phenotypes arising from loss of LRP10. This could include efforts to systematically test for additional lysosome proteins that depend on LRP10 for their efficient trafficking to lysosomes. Likewise, the reasons for selective vulnerability of microglia to LRP10 depletion warrant investigation. Generating conditional LRP10 knockout models to assess cell-type-specific effects on lysosomal function and neurodegeneration will be crucial. Additionally, investigating LRP10’s role in the trafficking of progranulin and prosaposin in human patient-derived cells and postmortem brain tissue could further strengthen understanding of the disease contributions of LRP10 insufficiency. Ultimately, targeting LRP10-related pathways may offer new therapeutic avenues for lysosomal dysfunction in neurodegenerative diseases.

## Materials and Methods

The Key Resource Table in the Supplemental Materials contains details of reagents and tools used in this study.

### Analysis of Gene Co-Expression

Genes of interest were entered as queries against “All Datasets” at SEEK (https://seek.princeton.edu/seek/)(Zhu *et al*, 2015).

### Culturing HeLa Cells

HeLa cells (kindly provided by Pietro De Camilli, Yale University) were cultured in Dulbecco’s Modified Eagle Medium (DMEM; Gibco, Thermo Fisher Scientific) supplemented with 10% fetal bovine serum (FBS; Gibco, Thermo Fisher Scientific), 100 U/mL penicillin, and 100 µg/mL streptomycin (Gibco, Thermo Fisher Scientific). Cells were maintained at 37°C in a humidified atmosphere containing 5% CO2. Cells were passaged every 2-3 days using 0.25% trypsin-EDTA (Gibco, Thermo Fisher Scientific) when they reached approximately 80% confluence. Details of stable cell lines are summerized in Key Resource Table. For plasmid DNA transfections, 1.8 - 2 x 10^5^ HeLa cells were plated on coverslips in a 24-well dish overnight. The next day, the transfection mix comprising 50 μl of OPTiMEM, 1 μl Lipofectamine 3000 (Invitrogen) and 0,5 μg of plasmid, was added to 500 μl of fresh new media, per well, and incubated for 24 hours. Transfections of RUSH plasmids were performed using Fugene 6 (Promega), seeding 1.5 x 10^5^ HeLa cells on coverslips in a 24-well dish overnight. The next day, the transfection mix made up of 50 μl of OPTiMEM, 1,5 μl Fugene 6 reagent and 0,5 μg of plasmid, was added to 500 μl of fresh new media, per well, and incubated for 24 hours. Cell culture and transfection protocols can be accessed at: https://doi.org/10.17504/protocols.io.8epv5rrq4g1b/v1and https://doi.org/10.17504/protocols.io.261ge55byg47/v1, respectively.

### CRISPR/Cas9 Genome Editing

Hela LRP10 KO cells were generated by transfection with px459-based Cas9 + sgRNA plasmids. The next day, 3µg/ml puromycin was added. 2 days later, cells were plated at single cell density in 96 well dishes. Clones were screened for loss of LRP10 by immunoblotting. A protocol can be accessed on protocols.io at: dx.doi.org/10.17504/protocols.io.dm6gp9bp5vzp/v.

### Lentivirus-Mediated Gene Expression

To generate stable cell lines via lentiviral transduction, HEK293FT cells were plated in a 6 well PDL coated plate and incubated overnight. The next day, cells were transfected with 100 μl of OPTiMEM, 3 μl of Fugene 6 and1 μg of a 1:1:1 ratio of psPAX2, pCMV-VSV-G and pLVx-HA-LRP10 plasmids. 24 hours before transduction, 1 x 10^6^ HeLa cells were seeded in a 6-well plate. The next day, media from HEK293FT cells was filtered, polybrene was added, and this was used to transduce the HeLa cells. Cells were selected with 3µg/ml puromycin the next day. A protocol can be accessed on protocols.io at: hyperlink in progress.

### Immunoblotting

3 x 10^5^ HeLa cells were seeded per well in a 6-well dish. The following day, cells were rinsed with PBS-1X and proteins were extracted by using cold lysis buffer (50 mM Tris-HCl pH 7.4, 150 mM NaCl, 1 % (v/v) Triton X-100, 1 mM EDTA supplemented with protease and phosphatase inhibitors). Cells were further centrifuged at 20,817 x g for 10 min at 4 °C and supernatants were recovered. Protein concentrations were measured with the Coomassie Plus Protein Assay Reagent (ThermoFisher Scientific). Proteins (15-30 ug) were diluted 1:1 with 2X β-mercaptoethanol sample buffer (80 mM Tris-HCl pH 6.8, 3.4 M Glycerol, 3 % (w/v) SDS, and 0.02 % (w/v) Bromophenol Blue supplemented with fresh 6 % (v/v) β-mercaptoethanol) and heated at 95 °C for 5 min. Protein lysates were loaded into 4-15 % miniPROTEAN TGX Stain-Free pre-cast gels (BioRad) using electrophoresis buffer (25 mM Tris Base, 192 mM Glycine, 0.1 % (w/v) SDS) and run at 250V for 25 min. Proteins were transferred onto 0.22 μm pore nitrocellulose membrane (Thermo Fisher Scientific) at 100 V for 60 min in transfer buffer (25 mM Tris Base, 192 mM Glycine, 20 % (v/v) Methanol). Membranes were then blocked in 5 % (w/v) non-fat dry milk diluted in a Tris-based saline solution with 0.1% of Tween 20 (TBST). Primary antibodies were added to the membranes in 5 % (w/v) BSA (Sigma-Aldrich) in TBST overnight (4 °C) at the indicated dilutions (Key Resource Table). Membranes were washed 3X for 5 minutes with TBST and incubated for 45 minutes with HRP-coupled secondary antibodies in TBST or 5 % (w/v) non-fat dry milk TBST. After 3 washes of 5 min in TBST, membranes were incubated with Pico ECL (Thermo Fisher Scientific) and the chemiluminescent signal was detected with a Chemidoc MP imager (Bio-Rad). A detailed protocol can be accessed on protocols.io at: https://doi.org/10.17504/protocols.io.bp2l6be9zgqe/v1.

### Immunofluorescence

HeLa cells were seeded on glass coverslips in 24-well plates at a density of 5 x 10^4^ cells per well. After 24 hours, cells were fixed with 8% paraformaldehyde (PFA) into the cell media (1:1 ratio) for 20 minutes at room temperature, followed by quenching of the reaction with 40mM PBS NH_4_Cl. Cells were blocked and permeabilized with PBS, 0.1% Saponin and 3% BSA for 15 min. Cells were then incubated overnight at 4°C with primary antibodies diluted in PBS, 0.1% Saponin and 3% BSA. After washing 3 times 5 min each with PBS, cells were incubated with appropriate secondary antibodies conjugated to fluorophores (e.g., Alexa Fluor 488, 568; Invitrogen, Thermo Fisher Scientific) for 1 hour at room temperature in the dark. Nuclei were counterstained with 4’,6-diamidino-2-phenylindole (DAPI; Invitrogen, Thermo Fisher Scientific). Coverslips were mounted on glass slides using ProLong Gold Antifade Mountant (Invitrogen, Thermo Fisher Scientific), and images were captured using a confocal microscope. A protocol can be accessed on protocols.io at: 10.17504/protocols.io.81wgbym7ovpk/v1.

### RUSH Assay

RUSH plasmids were used as previously described (Devireddy and Ferguson, 2021). Briefly, Hela cells were transfected with RUSH plasmids encoding an ER hook and mCherry as reporter. Cells were cultured as mentioned above. Following addition of 40 µM biotin, cells were fixed at indicated timepoints before immunostaining. A protocol can be accessed on protocols.io at: Hyperlink in Progress

### Lysosome Isolation

Lysosomes were magnetically isolated from HeLa cells by endocytic loading of cells with supermagnetic iron oxide nanoparticles (SPIONs) that were prepared as previously described (Amick et al., 2018; Hancock-Cerutti et al, 2022; Rodriguez-Paris et al.,1993). For cell homogenization, this current study employed a Dounce homogenizer (tight pestle) rather than the Isobiotec homogenizer. Briefly, HeLa cells 1.5×10^6^ were seeded in a 15 cm dish. The next day, the media was replaced with fresh media containing 10% SPION particles supplemented with 5mM HEPES (ph 7.4, 15630-080; Thermo Fisher Scientific) and left for 24 h. After two washes with cold PBS, cells were harvested in cold PBS and centrifuged at 4° for 10 min at 300g to remove nuclei and cell debris. The cells were resuspended in 1 ml/dish of ice-cold HB buffer (5 mM Tris base [AB02000-05000; American Bio], 250 mM sucrose [S0389; Sigma-Aldrich], and 1 mM EGTA pH 7.4 [E4378; Sigma-Aldrich]) supplemented with inhibitors (cOmplete mini EDTA-free protease inhibitor [11836170001; Roche] and PhosSTOP [4906837001; Roche]) and passed 50 times through a Dounce homogenizer (DWK Life Sciences Wheaton Dounce Tissue Grinders, 06-434) with a tight pestle to generate the total cell lysate which was next centrifuged at 4°C for 10 min at 800g to obtain the post-nuclear supernatant, a portion of which was kept. LS columns (130042401; Miltenyi Biotec) were washed once with 2.5 ml of HB buffer on a QuadroMACS separator (130-091-051; Miltenyi Biotec). The remainder of the cell lysate was applied to LS columns. The flow-through was collected and reapplied on the column. The columns were washed with 5 ml of HB buffer, removed from the magnetic rack, and the bound fraction was eluted into ultracentrifuge tubes using 3 ml of HB buffer. Samples were ultracentrifuged at 55,000g at 4°C for 10 min using a TLA-100.3 rotor in a Beckman-Coulter Ultracentrifuge Max Optima to concentrate samples and thereby generating the lysosomal pellet. The supernatant was removed, and the lysosome pellet was resuspended in ∼50 μl of HB buffer and prepared for immunoblotting. Protein concentration was determined using the BCA Protein Assay Kit (Pierce, Thermo Fisher Scientific). Lysosomes were subsequently analyzed by immunoblotting. More detailed protocols for the SPION preparation and purification of lysosomes with dextran conjugated SPIONs can be accessed on protocols.io at: dx.doi.org/10.17504/protocols.io.eq2lyn69pvx9/v1and dx.doi.org/10.17504/protocols.io.bp2l61dr1vqe/v1.

### LRP10 Mutant Mice

The mouse LRP10 KO was generated by the Yale Genome Editing Center. To this end, C57Bl/6J zygotes were electroporated with a pair of Cas9/sgRNA ribonucleoprotein complexes that targeted sequences in exon 1 (CCGGCACCCCTGTTCAACCC)and intron 1 (AGAGGAGGCCGAGGCTCGAC) respectively. This resulted in the deletion of a 128 bp fragment which contained the LRP10 start codon. Genotyping of LRP10 mutant mice was carried out by polymerase chain reaction on DNA from tail biopsies using oligonucleotide primers (GTCACGCCTGTTCCTCTCC and ACCTCCTAGAGGGGGAATCG) that yielded bands of 485 base pairs for the wildtype allele and 357 base pairs for the mutant allele.

### Perfusion, Vibratome Cutting, and Immunohistochemistry of Mouse Brains

Mice were anesthetized with isoflurane and transcardially perfused with ice-cold PBS followed by 4% PFA in PBS. Brains were dissected and post-fixed in 4% PFA at 4°C overnight. Brains were then rinsed in PBS and embedded in 20% sucrose solution overnight and then kept in 30% sucrose for storage. Coronal sections (40-50 µm) were cut using a vibratome (Leica VT1000S) and collected in cold PBS with sodium-azide. Free-floating sections were permeabilized with 0.3% Triton X-100 in PBS for 1 hour at room temperature and blocked with 5% normal donkey serum (NDS) in PBS for 2 hours at room temperature. Sections were incubated with primary antibodies diluted in PBS containing 1% NDS and 0.1% Triton X-100 overnight at 4°C. After washing with PBS, sections were incubated with appropriate fluorophore-conjugated secondary antibodies for 2 hours at room temperature in the dark. Sections were mounted on glass slides using Fluoromount-G (SouthernBiotech), and images were captured using a confocal microscope. A protocol can be accessed at: dx.doi.org<u>/10.17504/protocols.io.ewov1obeolr2/v1</u>.

### Confocal Imaging

Z-stack images were taken using a Nikon SoRa spinning disk confocal microscope (Apo TIRF 60x Oil NA 1.49), with sCMOS camera, in normal and SoRa mode. Zeiss LSM 900 scanning confocal microscope (plan-apochromat 63x/1.4 Oil DIC M27), with GaAsP PMTs detection, was used for better optical sectioning. Sequential scanning and low-laser intensity setting were used to avoid signal bleed-through.

### Data Analysis

Fluorescence images were analyzed using Fiji (version 2.16.0) and Icy software (version 2.0) ((Schindelin *et al*, 2012; de Chaumont *et al*, 2012). Colocalization, morphology, number of cells and intensity measurements were performed using Coloc2, Sholl, Analyse particles and Measure plugins in Fiji, respectively. 3D reconstruction analysis were performed using Icy. RUSH experiments were quantified by defining a whole cell mask and a Golgi mask and measuring both integrated density. Post-Golgi trafficking ratio was calculated by using the following formula (%) = [(Whole RUSH signal-GM130 RUSH signal) / Whole RUSH signal] x 100 for each cell. Quantitative data were obtained from at least three independent experiments and were expressed as mean ± standard error of mean (SEM). Pooled data from different replicate experiments were standardized as follows: “normalized value (x) = (x – sample mean)/sample standard deviation”. To avoid negative numbers, the lowest value of all replicates was subtracted from the totality of values in the dataset. Details of quantification tools are listed in the Key Resource Table.

### Data Representation and Statistical Analysis

Calculations were performed in Microsoft Excel (version 16.96.1). Statistical analyses and graphs generation were performed using GraphPad Prism (version 10). Comparisons between two groups were made using unpaired Student’s t-test using Welch’s correction, while multiple group comparisons were performed using one-way ANOVA followed by Tukey’s post hoc test. Mean ± S.E.M of at least three independent experiments are shown. P-values = ∗ p < 0.05; ∗∗ p < 0.01; ∗∗∗ p < 0.001; ∗∗∗∗ p < 0.0001; ns, not significant). Data from microscopy images were displayed as Superplots, a data visualization method that allows to emphasize both within-experiment variability and experimental reproducibility (Lord *et al*, 2020) Figures were generated using open source Inkscape software (version 1.2.1).

## Acknowledgements

This research was supported by grants from Aligning Science Across Parkinson’s disease (ASAP-000580) through the Michael J. Fox Foundation for Parkinson’s Research (MJFF), The Bluefield Foundation and The Parkinson’s Disease Foundation (PF-RCE-1946). Agnes Roczniak-Ferguson (Yale) provided valuable guidance and feedback. We are grateful to Suxia Bai and Timothy Nottoli at the Yale Genome Editing Center for generation of the LRP10 mutant mice and to Taylor Skibitcky for help with mouse genotyping. Agnes Roczniak-Ferguson provided technical and intellectual guidance throughout the project and contributed immensely to maintaining a productive laboratory environment. We thank Berrak Ugur (Yale) and Benjamin Johnson (Yale) for managing our ASAP project. The authors do not have any conflicts to declare.

