## Supplemental Data for "LRP10 promotes trafficking of progranulin and prosaposin to lysosomes"

#### Supplemental Figures

**A**

| Term | p-value | q-value |
|---|---|---|
| **Vacuolar part** | 1.40E-13 | 8.31E-14 |
| **Lysosomal lumen** | 1.40E-13 | 8.31E-14 |
| **Vacuole** | 3.57E-13 | 4.05E-13 |
| **Lysosome** | 4.42E-13 | 4.56E-13 |
| Phagocytic vesicle membrane | 9.05E-13 | 1.60E-11 |
| Integral to lumenal side of endoplasmic reticulum membrane | 1.54E-06 | 6.74E-06 |
| Endosomal part | 2.81E-06 | 7.36E-06 |
| Endosomal membrane | 1.45E-05 | 3.99E-05 |
| Endosome | 1.24E-04 | 1.43E-04 |
| ER to Golgi transport vesicle membrane | 1.24E-04 | 2.41E-04 |
| Integral to endoplasmic reticulum membrane | 2.01E-04 | 4.09E-04 |
| Cytoplasmic vesicle membrane | 2.33E-4 | 4.52E-04 |
| Integral to endoplasmic reticulum membrane | 8.48E-04 | 1.22E-03 |
| Cytoplasmic vesicle part | 1.86E-03 | 1.43E-03 |
| Membrane bounded vesicle | 1.86E-03 | 1.43E-03 |
| Transport vesicle | 1.86E-03 | 2.21E-03 |
| Lysosomal membrane | 1.96E-03 | 4.44E-03 |
| Early endosome | 4.56E-03 | 1.20E-02 |
| Cytoplasmic membrane bounded vesicle | 1.68E-02 | 1.50E-02 |
| Endoplasmic reticulum membrane | 4.12E-02 | 3.77E-02 |
| Golgi membrane | 4.81E-02 | 4.21E-02 |

**B**

| Term | p-value | q-value |
|---|---|---|
| **Vacuolar part** | 1.55E-07 | 2.35E-07 |
| **Vacuole** | 2.08E-07 | 2.35E-07 |
| **Lysosome** | 2.59E-07 | 2.41E-07 |
| Phagocytic vesicle membrane | 2.59E-07 | 3.21E-06 |
| Phagocytic vesicle | 3.26E-06 | 2.53E-05 |
| **Lysosomal lumen** | 1.77E-05 | 9.54E-05 |
| **Vacuolar membrane** | 1.67E-04 | 3.52E-04 |
| Endosomal part | 1.67E-04 | 3.84E-04 |
| Endocytic vesicle membrane | 1.67E-04 | 5.68E-04 |
| **Lysosomal membrane** | 4.52E-04 | 1.03E-03 |
| Membrane bounded vesicle | 1.63E-03 | 1.28E-03 |
| Endosome membrane | 1.63E-03 | 1.49E-03 |
| Endosome | 1.63E-03 | 1.73E-03 |
| Endocytic vesicle | 1.63E-03 | 2.03E-03 |
| Integral to lumenl side of endoplasmic reticulum membrane | 1.63E-03 | 2.30E-03 |
| Cytoplasmic vesicle part | 1.46E-02 | 2.11E-02 |
| Integral to endoplasmic reticulum membrane | 1.46E-02 | 2.79E-02 |
| Cytoplasmic membrane bounded vesicle | 4.84E-02 | 4.10E-02 |
| ER to Gogli transport vesicle membrane | 4.84E-02 | 4.10E-02 |
| Golgi membrane | 4.84E-02 | 4.10E-02 |
| Extracellular vesicular exosome | 8.13E-02 | 6.57E-02 |
| Transport vesicle | 8.13E-02 | 8.78E-02 |
| Gogli apparatus part | 8.22E-02 | 8.78E-02 |
| Cleavage furrow | 8.22E-02 | 1.27E-01 |
| Stress fiber | 8.22E-02 | 2.16E-01 |
| Proton transporting two sector ATPase complex | 8.22E-02 | 2.39E-01 |

**Figure S1. Gene ontology (GO) enrichment analyses of co-expression data**

(A and B) Tables showing results of enrichment of GO cellular components from the results of GRN (A) and LRP10 (B) SEEK co-expression analyses. Lysosome-related terms are displayed in blue and the P-value is adjusted for multiple hypothesis testing (Benjamini-Hochberg), q-value represents minimum false discovery rate. The top100 co-expressed genes were used for these analyses.

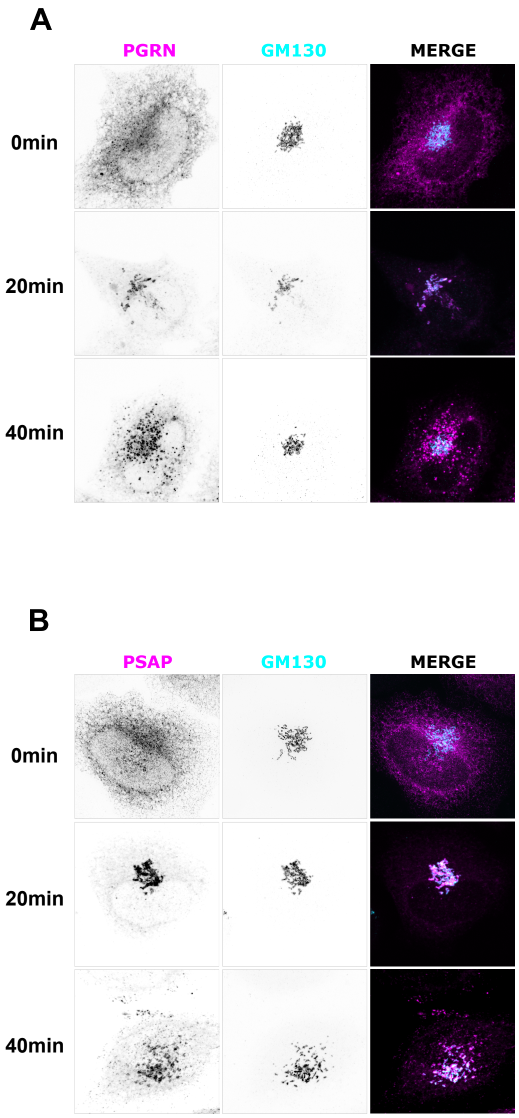


**Figure S2. Dynamic trafficking of PGRN and PSAP in the RUSH assay**

(A-B) Spinning disk confocal imaging of mCherry-RUSH-PGRN and mCherry-PSAP-RUSH expressed in wild-type Hela cells at t = 0, 20, 40 min and 60 min. In both experiments, cells were co-stained with an mCherry antibody for detection of progranulin or prosaposin and for GM130 to detect the Golgi apparatus.

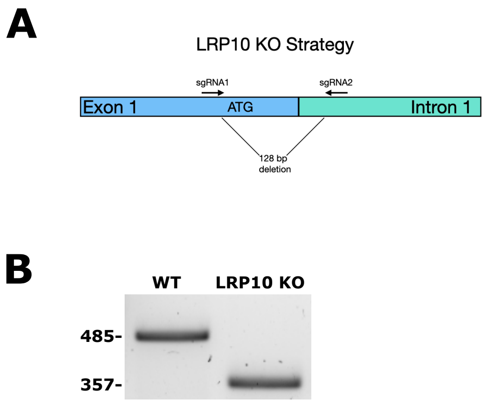


**Figure S3. Generation of LRP10 KO mice**

(A) Schematic diagram of the strategy for generating LRP10 KO mice. (B)Representative PCR gel showing the genotyping results for wild-type and knock-out LRP10 mice. DNA was extracted from mouse tail biopsies and amplified using primers specific for the LRP10 wild-type and knock-out alleles. The wild-type allele produces a band of 485 bp, while the knock-out allele generates a band of 357bp.

A

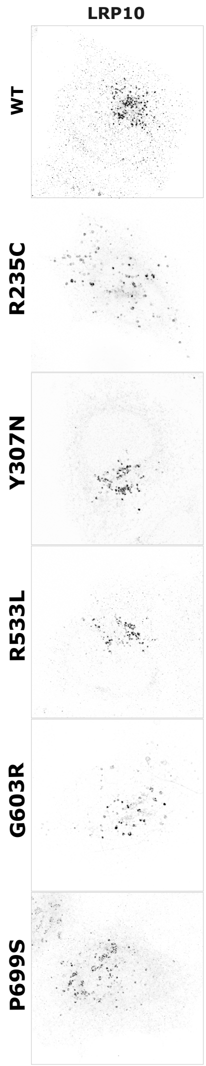


**Figure S4. Subcellular localization of disease-associated LRP10 mutant proteins**

(A) LRP10 knockout Hela cells were transfected with wildtype or the indicated mutant HA-LRP10 plasmids and processed for immunofluorescence detection of LRP10. Scale bar, 5 µm.

### Key Resource Table

| RESOURCE TYPE | RESOURCE NAME | SOURCE | IDENTIFIER | NEW/REUSE | RRID |
|---|---|---|---|---|---|
| **Antibody** | HA | Cell Signaling Technologies | HA-tag (6E2) | REUSE | AB_10691311 |
| **Antibody** | LRP10 | MRC PPU Reagents | DA058 | REUSE | AB_3678594 |
| **Antibody** | TGN46 human | Bio-Rad | AHP500G | REUSE | AB_323104 |
| **Antibody** | TGN46 mouse | Invitrogen | MA3-063 | REUSE | AB_325484 |
| **Antibody** | EEA1 | Termofisher | PA1-063A | REUSE | AB_2096819 |
| **Antibody** | Cathepsin D human | R&D | AF1014 | REUSE | AB_2087218 |
| **Antibody** | Cathepsin D mouse | R&D | AF1029 | REUSE | AB_2087094 |
| **Antibody** | Progranulin human | R&D | AF2420 | REUSE | AB_2114489 |
| **Antibody** | Progranulin mouse | R&D | AF2557 | REUSE | AB_2114504 |
| **Antibody** | P230 | Biosciences | 611280 | REUSE | AB_398808 |
| **Antibody** | Prosaposin human/Saposin C | Santa Cruz Biotechnology | sc-374118 | REUSE | AB_10915437 |
| **Antibody** | Prosaposin mouse | Abcam | ab300469 | REUSE | AB_3101805 |
| **Antibody** | LAMP1 human | Hybridoma/J. Thomas August | H4A3 | REUSE | AB_2296838 |
| **Antibody** | LAMP1 human | Cell Signaling Technologies | D2D11 | REUSE | AB_2687579 |
| **Antibody** | LAMP1 mouse | Hybridoma/J. Thomas August | 1D4B | REUSE | AB_528127 |
| **Antibody** | GBA | R&D | MAB7410 | REUSE | AB_2938856 |
| **Antibody** | Granulin 2/3 | Sigma-Aldrich | HPA008763 | REUSE | AB_1850339 |
| **Antibody** | PDI | Cell Signaling Technologies | 2446S | REUSE | AB_2298935 |
| **Antibody** | GM130 | BD Biosciences | 610822 | REUSE | AB_398141 |
| **Antibody** | RFP | Chromostek | 5f8 | REUSE | AB_2336064 |
| **Antibody** | IBA1 | Fujifilm Wako's | 019-19741 | REUSE | AB_839504 |
| **Antibody** | Beta-III-tubulin | Aves labs | TUJ | REUSE | AB_2313564 |
| **Antibody** | Actin | Santa Cruz Biotechnology | sc-8432 | REUSE | AB_626630 |
| **Antibody** | Rabbit IgG (HRP) | Cell Signaling Technologies | 7074S | REUSE | AB_2099233 |
| **Antibody** | Mouse IgG (HRP) | Cell Signaling Technologies | 7076S | REUSE | AB_330924 |
| **Antibody** | Goat IgG (HRP) | Cell Signaling Technologies | 91196S | REUSE | AB_2940774 |
| **Antibody** | Biotin (HRP) | Cell Signaling Technologies | 7075S | REUSE | AB_10696897 |
| **Antibody** | AlexaFluor 568 anti-mouse | Invitrogen | A10037 | REUSE | AB_11180865 |
| **Antibody** | AlexaFluor 488 anti-goat | Invitrogen | A11055 | REUSE | AB_10679538 |
| **Antibody** | AlexaFluor 488 anti-rabbit | Invitrogen | A11008 | REUSE | AB_143165 |
| **Antibody** | AlexaFluor 568 anti-rat | Invitrogen | A11077 | REUSE | AB_2534121 |
| **Antibody** | AlexaFluor 647 anti-rabbit | Invitrogen | A31573 | REUSE | AB_2536183 |
| **Antibody** | AlexaFluor 488 anti-sheep | Invitrogen | A11015 | REUSE | AB_141362 |
| **Antibody** | AlexaFluor 568 anti-mouse | Invitrogen | A11004 | REUSE | AB_2534072 |
| **Antibody** | AlexaFluor 568 anti-rabbit | Invitrogen | A11011 | REUSE | AB_143157 |
| **Antibody** | AlexaFluor 568 anti-chicken | Invitrogen | A11041 | REUSE | AB_2534098 |
| **Recombinant DNA** | pLVX-Puro-hLRP10 | This paper | | NEW | |
| **Recombinant DNA** | STR-KDEL_PSAP-SBP-mCherry | Devireddy and Ferguson, 2021 | Addgene 180440 | REUSE | Addgene_180440 |
| **Recombinant DNA** | STR-KDEL_SBP-mCherry-PGRN | Devireddy and Ferguson, 2021 | Addgene 180439 | REUSE | Addgene_180439 |
| **Recombinant DNA** | SS-HA-LRP10-WT | This paper | | NEW | |
| **Recombinant DNA** | SS-HA-LRP10-R235C | This paper | | NEW | |
| **Recombinant DNA** | SS-HA-LRP10-Y307N | This paper | | NEW | |

| RESOURCE TYPE | RESOURCE NAME | SOURCE | IDENTIFIER | NEW/REUSE | RRID |
|---|---|---|---|---|---|
| **Recombinant DNA** | SS-HA-LRP10-R533L | This paper | | NEW | |
| **Recombinant DNA** | SS-HA-LRP10-G603R | This paper | | NEW | |
| **Recombinant DNA** | SS-HA-LRP10-P699S | This paper | | NEW | |
| **Experimental model: Cell line** | Hela M WT | Pietro De Camilli, Yale University | | REUSE | |
| **Experimental model: Cell line** | Hela M | LRP10 KO | This paper | NEW | |
| **Chemical, peptide, or recombinant protein** | DMEM | Thermo Fisher Scientific | 11965-092 | REUSE | |
| **Chemical, peptide, or recombinant protein** | FBS | Thermo Fisher Scientific | 16140-071 | REUSE | |
| **Chemical, peptide, or recombinant protein** | PBS | Thermo Fisher Scientific | 10010023 | REUSE | |
| **Chemical, peptide, or recombinant protein** | Cell Stripper | Corning | 25056CI | REUSE | |
| **Chemical, peptide, or recombinant protein** | Penicillin/Streptomicin (10,000 U/mL) | Thermo Fisher Scientific | 15140122 | REUSE | |
| **Chemical, peptide, or recombinant protein** | Puromycin | Thermo Fisher Scientific | A11138-03 | REUSE | |
| **Chemical, peptide, or recombinant protein** | Opti-Mem | Thermo Fisher Scientific | 31985062 | REUSE | |
| **Chemical, peptide, or recombinant protein** | Lipofectamine 2000 | Invitrogen | 11668019 | REUSE | |
| **Chemical, peptide, or recombinant protein** | Lipofectamine CRISPRMAX | Invitrogen | CMAX00003 | REUSE | |
| **Chemical, peptide, or recombinant protein** | Fugene HD | Promega | E2311 | REUSE | |
| **Chemical, peptide, or recombinant protein** | Potassium Phosphate Monobasic | J.T. Baker | Jan-46 | REUSE | |
| **Chemical, peptide, or recombinant protein** | Sodium Phosphate Dibasic | J.T. Baker | May-28 | REUSE | |
| **Chemical, peptide, or recombinant protein** | Glycine | American Bio | AB00730-05000 | REUSE | |
| **Chemical, peptide, or recombinant protein** | Tris | American Bio | AB02000-05000 | REUSE | |
| **Chemical, peptide, or recombinant protein** | NaCl | Sigma-Aldrich | May-24 | REUSE | |
| **Chemical, peptide, or recombinant protein** | Hydrochloric Acid | J.T. Baker | 9535 | REUSE | |
| **Chemical, peptide, or recombinant protein** | SDS | American Bio | AB01920-00500 | REUSE | |
| **Chemical, peptide, or recombinant protein** | EDTA | Sigma-Aldrich | 3690 | REUSE | |
| **Chemical, peptide, or recombinant protein** | Triton X-100 | Sigma-Aldrich | X100 | REUSE | |
| **Chemical, peptide, or recombinant protein** | Tween-20 | Sigma-Aldrich | P7949 | REUSE | |
| **Chemical, peptide, or recombinant protein** | Glycerol | American Bio | AB00751 | REUSE | |
| **Chemical, peptide, or recombinant protein** | Bromphenol Blue | Sigma-Aldrich | B5525 | REUSE | |
| **Chemical, peptide, or recombinant protein** | B-mercaptoethanol | Sigma-Aldrich | M3148 | REUSE | |
| **Chemical, peptide, or recombinant protein** | Sucrose | Sigma-Aldrich | S0389 | REUSE | |
| **Chemical, peptide, or recombinant protein** | EGTA | Sigma-Aldrich | E4378 | REUSE | |

| RESOURCE TYPE | RESOURCE NAME | SOURCE | IDENTIFIER | NEW/REUSE | RRID |
|---|---|---|---|---|---|
| **Chemical, peptide, or recombinant protein** | HEPES (pH 7.4) | Thermo Fisher Scientific | 15630-080 | REUSE | |
| **Chemical, peptide, or recombinant protein** | DMSO | Sigma-Aldrich | D2650 | REUSE | |
| **Chemical, peptide, or recombinant protein** | COmplete mini EDTA Free | Roche | 11836170001 | REUSE | |
| **Chemical, peptide, or recombinant protein** | PhosSTOP | Roche | 4906837001 | REUSE | |
| **Chemical, peptide, or recombinant protein** | Coomassie Plus Protein Assay Reagent | Thermo Fisher Scientific | 23236 | REUSE | |
| **Chemical, peptide, or recombinant protein** | PAGEruler Plus Prestained Protein Ladder | Thermo Fisher Scientific | 26620 | REUSE | |
| **Chemical, peptide, or recombinant protein** | Biotin Protein Ladder | Cell Signaling | 7727L | REUSE | |
| **Chemical, peptide, or recombinant protein** | 4-15% MiniPROTEAN 10-well | Biorad | 4568084g | REUSE | |
| **Chemical, peptide, or recombinant protein** | BSA | Sigma-Aldrich | A9647 | REUSE | |
| **Chemical, peptide, or recombinant protein** | Non-Fat Dry Milk Omniblock | American Bio | AB10109-01000 | REUSE | |
| **Chemical, peptide, or recombinant protein** | 0.45 um Nitrocellulose Membrane | Thermo Fisher Scientific | 1620115 | REUSE | |
| **Chemical, peptide, or recombinant protein** | Whatman Filter Paper | VWR | 28298-020 | REUSE | |
| **Chemical, peptide, or recombinant protein** | SuperSignal West Pico PLUS Chemiluminescence Substrate | Thermo Fisher Scientific | 34580 | REUSE | |
| **Chemical, peptide, or recombinant protein** | SuperSignal West Femto Maximum Sensitivity Substrate | Thermo Fisher Scientific | 34095 | REUSE | |
| **Chemical, peptide, or recombinant protein** | Methanol | Sigma-Aldrich | 179337-4L-PB | REUSE | |
| **Chemical, peptide, or recombinant protein** | Ethanol | Decon Laboratories | 2716 | REUSE | |
| **Chemical, peptide, or recombinant protein** | Ampicillin | Sigma-Aldrich | A0166 | REUSE | |
| **Chemical, peptide, or recombinant protein** | LB + Ampicillin (100 µg/mL) | Recombinant Technologies | 760100 | REUSE | |
| **Chemical, peptide, or recombinant protein** | Iron (II) Chloride | Sigma-Aldrich | 220299 | REUSE | |
| **Chemical, peptide, or recombinant protein** | Iron (III) Chloride | Sigma-Aldrich | 157740 | REUSE | |
| **Chemical, peptide, or recombinant protein** | Ammonium hydroxide (30%) | Sigma-Aldrich | 320145 | REUSE | |
| **Chemical, peptide, or recombinant protein** | Dextran | Sigma-Aldrich | D1662 | REUSE | |
| **Chemical, peptide, or recombinant protein** | Snakeskin dialysis tubing (10,000 Mol Wt) | Thermo Fisher Scientific | 68100, 10,000 | REUSE | |
| **Chemical, peptide, or recombinant protein** | LS Columns | Miltenyi Biotec | 130-042-401 | REUSE | |
| **Chemical, peptide, or recombinant protein** | QuadroMACS Separator | Miltenyi Biotec | 130-091-051 | REUSE | |
| **Chemical, peptide, or recombinant protein** | Saponin Quilajja sp. | Sigma-Aldrich | S4521 | REUSE | |
| **Chemical, peptide, or recombinant protein** | Paraformaldehyde | Electron Microscopy Sciences | 19202 | REUSE | |
| **Chemical, peptide, or recombinant protein** | Sodium dihydrogen phosphate monohydrate | J.T. Baker | 3818 | REUSE | |

| RESOURCE TYPE | RESOURCE NAME | SOURCE | IDENTIFIER | NEW/REUSE | RRID |
|---|---|---|---|---|---|
| **Chemical, peptide, or recombinant protein** | Sodium phosphate, dibasic, anhydrous | J.T. Baker | 3828 | REUSE | |
| **Chemical, peptide, or recombinant protein** | ProLong™ Gold Antifade Mountant with DNA Stain DAPI | Thermo Fisher Scientific | P36935 | REUSE | |
| **Chemical, peptide, or recombinant protein** | Fisherbrand™ Superfrost™ Disposable Microscope Slides | Thermo Fisher Scientific | 12-550-143 | REUSE | |
| **Chemical, peptide, or recombinant protein** | Microscope Cover Slips (12 mm) | Carolina Biological Supply | 633029 | REUSE | |
| **Chemical, peptide, or recombinant protein** | Q5 High-Fidelity 2X Master Mix | NEB | M0492S | REUSE | |
| **Chemical, peptide, or recombinant protein** | HIFI DNA Assembly Master Mix | NEB | E2621L | REUSE | |
| **Chemical, peptide, or recombinant protein** | One-Shot STABL3 | Invitrogen | C7373-03 | REUSE | |
| **Software/Code** | Prism 10 | Graphpad | SCR_002798 | REUSE | |
| **Software/Code** | FIJI 2.14.0/1.54f | | SCR_002285 | REUSE | |
| **Software/Code** | ChimeraX 1.7.1 | | SCR_015872 | REUSE | |
